# Phytoplankton trigger the production of cryptic metabolites in the marine actinobacteria *Salinispora tropica*

**DOI:** 10.1101/2020.05.18.103358

**Authors:** Audam Chhun, Despoina Sousoni, Maria del Mar Aguiló-Ferretjans, Lijiang Song, Christophe Corre, Joseph A. Christie-Oleza

## Abstract

Bacteria from the Actinomycete family are a remarkable source of natural products with pharmaceutical potential. The discovery of novel molecules from these organisms is, however, hindered because most of the biosynthetic gene clusters (BGCs) encoding these secondary metabolites are cryptic or silent and are referred to as orphan BGCs. While co-culture has proven to be a promising approach to unlock the biosynthetic potential of many microorganisms by activating the expression of these orphan BGCs, it still remains an underexplored technique. The marine actinobacteria *Salinispora tropica*, for instance, produces valuable compounds such as the anti-cancer molecule salinosporamide A but half of its putative BGCs are still orphan. Although previous studies have looked into using marine heterotrophs to induce orphan BGCs in *Salinispora*, the potential impact of co-culturing marine phototrophs with *Salinispora* has yet to be investigated. Following the observation of clear antimicrobial phenotype of the actinobacterium on a range of phytoplanktonic organisms, we here report the discovery of novel cryptic secondary metabolites produced by *S. tropica* in response to its co-culture with photosynthetic primary producers. An approach combining metabolomics and proteomics revealed that the photosynthate released by phytoplankton influences the biosynthetic capacities of *S. tropica* with both production of new molecules and the activation of orphan BGCs. Our work pioneers the use of phototrophs as a promising strategy to accelerate the discovery of novel natural products from actinobacteria.

**Importance:** The alarming increase of antimicrobial resistance has generated an enormous interest in the discovery of novel active compounds. The isolation of new microbes to untap novel natural products is currently hampered because most biosynthetic gene clusters (BGC) encoded by these microorganisms are not expressed under standard laboratory conditions, *i.e.* mono-cultures. Here we show that co-culturing can be an easy way for triggering silent BGC. By combining state-of-the-art metabolomics and high-throughput proteomics, we characterized the activation of cryptic metabolites and silent biosynthetic gene clusters in the marine actinobacteria *Salinispora tropica* by the presence of phytoplankton photosynthate. We further suggest a mechanistic understanding of the antimicrobial effect this actinobacterium has on a broad range of prokaryotic and eukaryotic phytoplankton species and reveal a promising candidate for antibiotic production.

## Introduction

Soil actinomycetes are a rich source of drug-like natural products, to which we owe up to 70% of all microbial antibiotics used today (Bérdy, 2005). Identification of novel secondary metabolites from this extensively studied family has, however, stalled over the last few decades as a result of the recurring rediscovery of already known compounds. This has led in recent years to a thriving interest for the study of new microorganisms, with the rational that ecologically distinct microorganisms produce equally distinct secondary metabolites (Molinski et *al.*, 2009; Wilson and Brimble, 2009). For instance, the heterotrophic bacteria *Salinispora* drew particular attention when discovered, as it was the first obligate marine actinomycete described (Jensen *et al*., 1991; Mincer *et al*., 2002, Jensen & Mafnas, 2006) and has since proven to be an important source of new natural products for the pharmaceutical industry (Maldonado *et al.*, 2005; Feling *et al.*, 2003; Buchanan *et al*., 2005; Asolkar *et al*., 2010). Despite the increasing number of novel strains identified with promising biosynthetic capacities, many hurdles in natural product discovery remain. Most of these microbial secondary metabolites are encoded by groups of colocalized genes, called biosynthetic gene clusters (BGCs), which are now more easily identified because of the improvement in sequencing technologies and bioinformatic tools (Medema *et al*., 2011). The majority of these discovered BGCs, however, have yet to be linked to their products and are called orphan BGCs. They are generally considered to be either silent - because of a low level of expression or inactivation of their biosynthetic genes - or the metabolites they produce are cryptic - difficult to detect and isolate (Reen *et al*., 2015; Rutledge and Challis, 2015). The observation of numerous orphan BGCs in genome-sequenced microorganisms has resulted in a growing interest in developing biological or chemical means to activate such clusters (Abdelmohsen *et al*., 2015; Onaka, 2017). One of the simplest and most efficient methods described in the literature relies on co-cultivation of different microbes to elicit novel natural product biosynthesis (Slattery *et al*., 2001; Bertrand *et al*., 2014).

The genome of the marine actinomycete *Salinispora tropica* comprises at least 20 putative BGCs of which 11 are orphan (**Table 1**, Penn *et al.*, 2009, Udwary *et al*., 2007). Recent studies have shown that *Salinispora* co-inoculated with various marine heterotrophs could produce one or several antimicrobial compounds, which remain uncharacterized as traditional analytical chemistry methods did not allow their identification and no candidate BGC was proposed (Patin *et al*., 2016; Patin *et al*., 2018). While co-culturing appears to be a promising mean to activate orphan BGCs in *Salinispora*, it remains an underexplored technique to unravel the biosynthetic potential of this genus. Additionally, little has been done to establish the BGCs that are activated under such culturing conditions. Combining metabolomics with proteomics analyses has proven successful in linking novel compounds to active orphan BGCs in several *Streptomyces* species, but has not yet been applied to the genus *Salinispora* (Schley *et al*., 2006; Gubbens *et al*., 2014; Owens *et al.*, 2014).

**Table 1.**
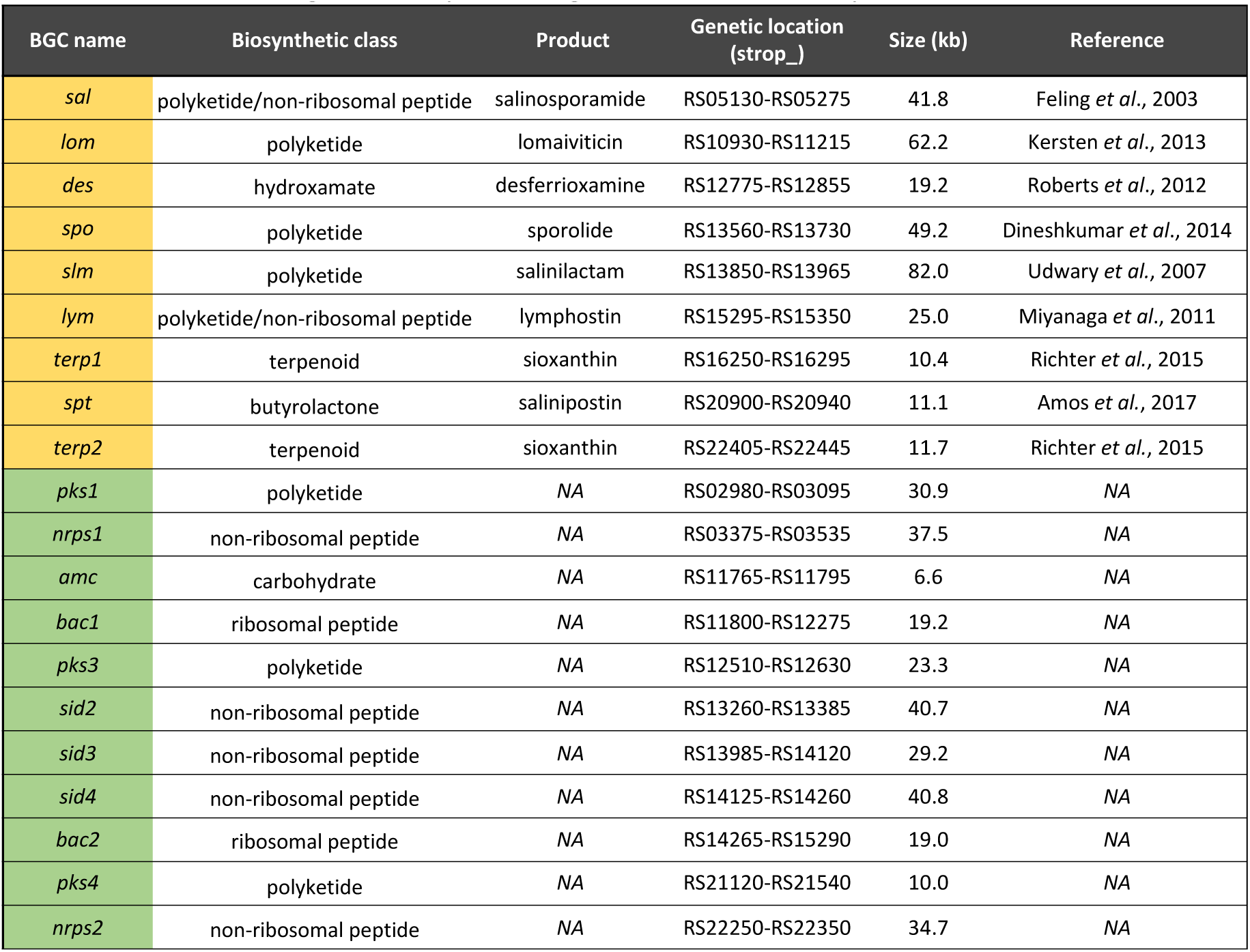
Biosynthetic gene clusters of *Salinispora tropica* CNB-440. Table shows the characterized (in orange) and orphan (in green) BGCs of *S. tropica* CNB-440.

Here we report the discovery of novel cryptic secondary metabolites produced by *S. tropica* CNB-440. By using an approach combining metabolomics and proteomics, we investigated how marine microbial phototrophs, and their photosynthate, induce the production of new metabolites and activate the expression of orphan BGCs in *S. tropica*. This strategy confirms microbial interactions as a promising and simple approach for future discovery of novel natural products.

## Material and methods

### 1. Culture conditions and cell abundance monitoring

#### 1.1. Strains and growth media

Axenic marine phototrophs *Synechococcus sp.* WH7803, *Emiliania huxleyi* RCC1242 and *Phaeodactylum tricornutum* CCAP1055/1 were routinely grown in Artificial Seawater (ASW, Wilson *et al*., 1996), K-media (Probert and Houdan, 2004), and F/2 media (Guillard *et al*., 1975), respectively. Cultures were set-up in Falcon 25 cm^2^ rectangular culture flasks with vented caps containing 20 ml of media and incubated at a constant light intensity of 10 μmol photons m^-2^ s^-1^, at 22 °C with orbital shaking (140 rpm). *Salinispora tropica* CNB-440 was grown in marine broth (MB, Difco), and incubated at 30 °C with orbital shaking (220 rpm). The *S. tropica* mutants *salA*^-^ and *salL*^-^ were generously provided by the Moore Laboratory, USA (Eustáquio *et al.*, 2008; Eustáquio *et al.*, 2009).

#### 1.2. Co-culture setup

*Salinispora* cells were grown to late exponential phase in 10 ml of MB before washing them three times with sterile mineral media, as appropriate for each phototroph, and finally resuspending the washed cell pellet in 10 ml of mineral media. Exponentially growing axenic phototroph cells and the washed *Salinispora* were co-inoculated in fresh media to a concentration of 10% (v/v) and 20% (v/v), respectively. *Salinispora* cells were also washed and resuspended in a conditioned *Synechococcus* supernatant (SUPSYN), when required for the metabolomic and proteomic analyses. To obtain the conditioned supernatant, *Synechococcus* cultures were incubated for 35 days as described above before centrifugation (4000 x *g* for 10 min at room temperature) and further filtration through 0.22 μm pore size filters to remove cells and particulate organic matter. Washed *Salinispora* cells were used to inoculate SUPSYN and MB, and cultures were incubated at 22°C with shaking (140 rpm) and a light intensity of 10 μmol photons m^-2^ s^-1^. For the physically separated *Synechococcus*-*Salinispora* co-cultures using the porous filters, cells were grown in 24 mm transwell with 0.4 μm pore polycarbonate membrane inserts (Corning). *Synechococcus* cells were inoculated in the well to a concentration of 20% (v/v) and *Salinispora* in the insert to a concentration of 55% (v/v).

#### 1.3. Flow cytometry

Phototroph cell abundance was monitored using their autofluorescence by flow cytometry using a LSR Fortessa Flow Cytometer (BD) instrument, and the BD FACSDiva acquisition software (BD). Cells were detected and gated using ex. 488 nm – em. 710/50 nm at voltage 370 V, and ex. 640 nm – em. filter 670/14 nm at voltage 480 V. To remove any *Salinispora* cell aggregates that would block the flow cell, samples were pre-filtered through a sterile mesh with pore size of 35 μm (Corning) prior to analysis.

### 2. Metabolomic analysis

#### 2.1. Sample preparation

The culture supernatants were analyzed by non-targeted metabolomic using either raw or concentrated supernatants. Raw supernatants were collected by sampling 200 μl of 0.22 μm-filtered culture milieu, prior to being mixed with an equal volume of HPLC-grade methanol. For concentrating the supernatant, cells from 10 to 100 mL of cultures were removed by centrifugation (4,000 x *g* for 15 min) followed by a filtering step using 0.22 μm vacuum filter bottle system (Corning). Pre-purification of the compounds of interest from the supernatants was carried out by solid phase extraction using C18-silica. Using a 90:10 A/B mobile phase (where A is water with 0.1% formic acid and B is methanol with 0.1% formic acid) the undesired polar molecules and salts passed through the silica while the compounds of interest were retained and later collected following elution with a 10:90 A/B mobile phase. The obtained fractions were dried under reduced pressure at 40 °C (in a speed-vac) and resuspended in 1-3 mL of 50:50 HPLC-grade methanol/water solution. All samples were stored in snap-seal amber glass vials (Thames Restek) and kept at -20 °C until analysis.

#### 2.2. Low-resolution LC-MS

Metabolites present in the cultures were routinely analyzed by reversed-phase liquid chromatography. A Dionex UltiMate 3000 HPLC (ThermoScientific) coupled with an amaZon SL Ion Trap MS (Bruker) was used. A Zorbax Eclipse Plus C18 column with dimensions 4.6 mm x 150 mm, 5 μm particle size (Agilent Technologies) was employed for metabolite separation with a linear gradient of 95:5 A/B to 30:70 A/B over 5 minutes, followed by second linear gradient to 20:80 A/B over 10 minutes with a flow rate of 1 ml min^-1^ (Mobile phase A: water with 0.1% formic acid, B: methanol with 0.1% formic acid). The mass spectrometer was operated in positive ion mode with a 100-1000 *m/z* scan range. The injected volume was 10 μL at a temperature of 25 °C. Data was processed with the Bruker Compass DataAnalysis software version 4.2 (Bruker).

#### 2.3. High-resolution LC-MS

To acquire molecular formulae information, samples were analyzed using an Ultra-high resolution MaXis II Q-TOF mass spectrometer equipped with electrospray source coupled with Dionex 3000RS UHPLC was employed (Bruker). A reverse phase C18 column (Agilent Zorbax, 100×2.1 mm, 1.8 μm) and a guard column (Agilent C18, 10×2.1 mm, 1.8 μm) were used for separation applying a linear gradient of 95:5 A/B to 0:100 A/B over 20 minutes (Mobile phase A: water with 0.1% formic acid, B: acetonitrile with 0.1% formic acid). The injected volume was 2 μL, and the flow rate was 0.2 ml min^-1^. At the beginning of each run, 7.5 μl of 10 mM of sodium formate solution was injected for internal calibration. The mass spectrometer was operated in positive ion mode with a 50-2500 *m/z* scan range. MS/MS data was acquired for the three most intense peaks in each scan.

### 3. Proteomic analysis

#### 3.1. Preparation of cellular proteome samples

Cultures were set up as described above and incubated for 5 days after which cells were collected by centrifuging 10 mL of culture at 4,000 x *g* for 10 min at 4 °C. Cell pellets were placed on dry ice before storing at -20 °C until further processing. The cell pellets were resuspended in 200 μL 1x NuPAGE lithium dodecyl sulfate (LDS) sample buffer (ThermoFischer Scientific), supplemented with 1% ß-mercaptoethanol. Cell pellets were lysed by bead beating (2×45 sec and 1×30 sec at 6.0 m/s) and sonication (5 min), followed by three successive 5-min incubations at 95 °C with short vortex steps in between. Cell lysates containing all proteins were loaded on an SDS-PAGE precast Tris-Bis NuPAGE gel (Invitrogen), using MOPS solution (Invitrogen) as the running buffer. Protein migration in the SDS-PAGE gel was performed for 5 min at 200 V, to allow removal of contaminants and purification of the polypeptides. The resulting gel was stained using SimplyBlue SafeStain (Invitrogen) to visualize the cellular proteome. The gel bands containing the cellular proteome were excised and stored at -20 °C until further processing.

#### 3.2. Trypsin in-gel digestion and nano LC-MS/MS analysis

Polyacrylamide gel bands were destained and standard in-gel reduction and alkylation were performed using dithiothreitol and iodoacetamide, respectively, after which proteins were in-gel digested overnight with 2.5 ng μL^-1^ trypsin (Christie-Oleza and Armengaud, 2010). The resulting peptide mixture was extracted by sonication of the gel slices in a solution of 5% formic acid in 25% acetonitrile, and finally concentrated at 40 °C in a speed-vac. For mass spectrometry analysis, peptides were resuspended in a solution of 0.05% trifluoroacetic acid in 2.5% acetonitrile prior to filtering using a 0.22 μm cellulose acetate spin column. Samples were analyzed by nanoLC-ESI-MS/MS with an Ultimate 3000 LC system (Dionex-LC Packings) coupled to an Orbitrap Fusion mass spectrometer (Thermo Scientific) using a 60 min LC separation on a 25 cm column and settings as previously specified (Christie-Oleza *et al*., 2015).

#### 3.3. Proteomic data analysis

Raw mass spectral files were processed for protein identification and quantification using the software MaxQuant (version 1.5.5.1; Cox and Mann, 2008) and the UniProt database of *S. tropica* CNB-440 (UP000000235). Quantification and normalization of spectral counts was done using the Label-Free Quantification (LFQ) method (Cox *et al*., 2014). Samples were matched between runs for peptide identification and other parameters were set by default. Data processing was completed using the software Perseus (version 1.5.5.3). Proteins were filtered by removing decoy and contaminants and were considered valid when present in at least two replicates for one condition. The relative abundance of each protein was calculated using protein intensities transformed to a logarithmic scale with base 2 and normalized to protein size. Variations in protein expression were assessed with a two-sample T-test, with a false discovery rate (FDR) *q* below 0.05 and a log(2) fold change above 2 (**Supplementary File S1**).

## Results

### *Salinispora tropica* has antimicrobial activity on a diverse range of marine phototrophs

Unlike other heterotrophs, which usually enhance the growth of phototrophic organisms when in co-culture (*e.g.* Christie-Oleza *et al*., 2017; Sher *et al*., 2011), *S. tropica* showed a clear antimicrobial activity on marine phytoplankton (**Fig. 1, A**). All three phototrophic model species tested, namely the cyanobacteria *Synechococcus* sp. WH7803, the coccolithophore *Emiliana huxleyi* and the diatom *Phaeodactylum tricornutum*, showed a strong decline in the presence of *S. tropica*, being especially remarkable for the two former species (**Fig. 1, A**). While also affected, the diatom *P. tricornutum* was not killed by *S. tropica* but, instead, its cells densities were significantly maintained one order of magnitude lower than when incubated axenically.

**Figure 1.**
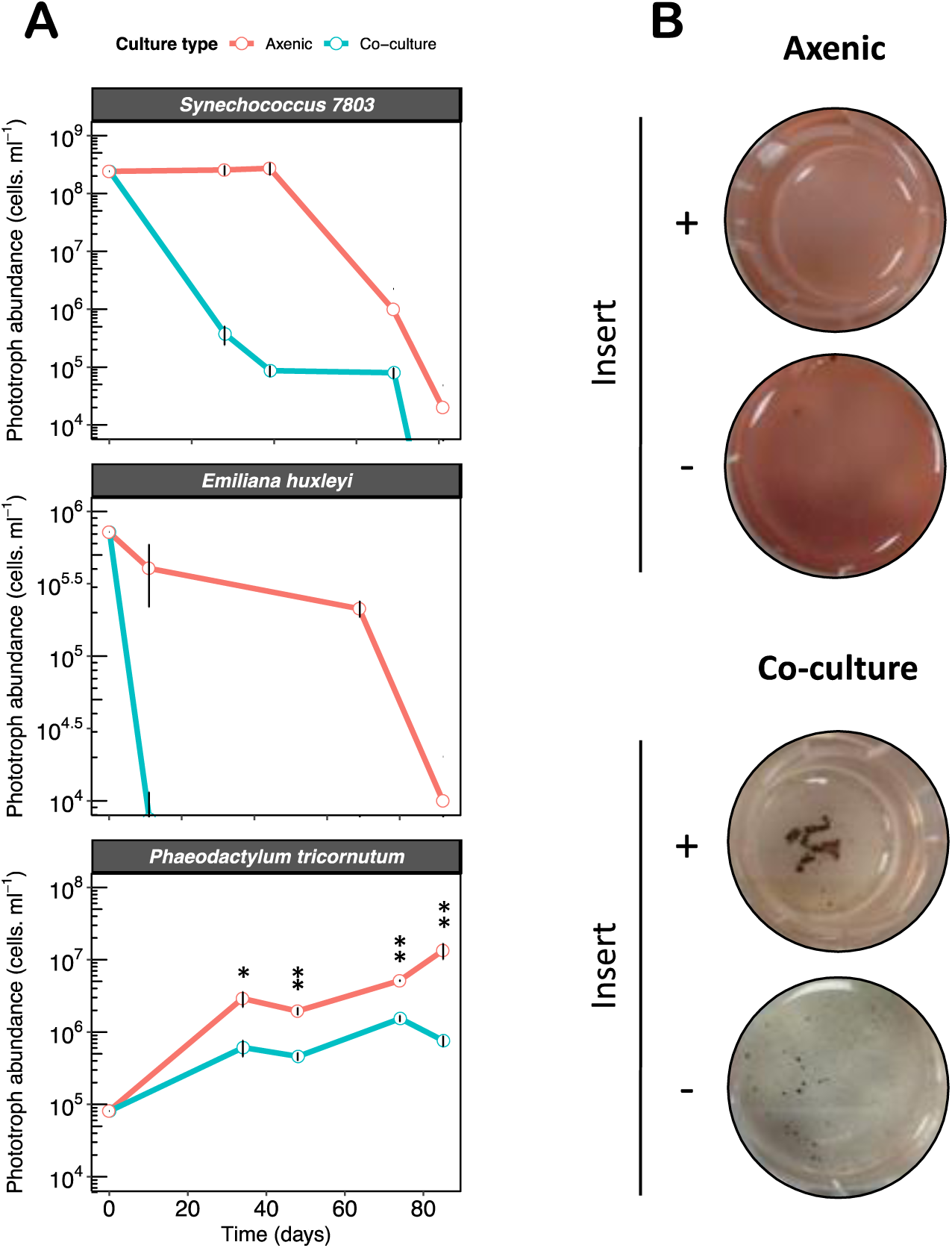
*Salinispora tropica* inhibits the growth of marine phototrophs *via* the secretion of an antimicrobial molecule. **(A)** *S. tropica* inhibits marine phototrophs growth in co-culture. Cultures of three marine phototrophs grown axenically (red lines) and in co-culture with *Salinispora tropica* (blue lines). Graph shows mean ± standard deviation of three biological replicates. Statistically significant cell abundances are indicated (T-test, significant * at *p*-value < 0.05 and ** at *p*-value < 0.01). **(B)** *Synechococcus* growth inhibition by *S. tropica* mediated by a diffusible molecule. The cyanobacterium was grown axenically and in co-culture with *S. tropica*, separated by a 0.4 µm pore membrane insert. Photographs of representative cultures of three biological replicates are shown, 7 days after inoculation. Red pigmentation is characteristic of healthy *Synechococcus* cells, while cell bleaching indicates cell death.

We were therefore interested in characterizing the nature of this inhibition. While other *Salinispora* species, such as *Salinispora arenicola*, are known to biosynthesize antibiotic molecules (Asolkar *et al*., 2010), no antimicrobial compound has yet been characterized in *S. tropica* CNB-440. Previous studies have shown, however, that *S. tropica* is able to outcompete other heterotrophs in co-culture by secreting siderophores leading to iron depletion (Patin *et al.*, 2016). To evaluate whether iron sequestration could explain the negative interactions observed in the present phototroph-*Salinispora* system, we supplemented the co-cultures with increasing concentrations of iron (**Supplementary Fig. S1**). The results obtained suggest that the antimicrobial phenotype was not due to siderophore activity, as saturating amount of iron could not rescue the growth of the phototrophs.

We then hypothesized that a yet unknown antimicrobial compound, to which our photosynthetic microorganisms are sensitive to, could be produced by *S. tropica.* To test this assumption, we setup co-cultures in which *S. tropica* and *Synechococcus* were physically separated by a porous filter, preventing direct cell-to-cell interactions while allowing the diffusion of small molecules (**Fig. 1, B**). *S. tropica* was still able to impair *Synechococcus* proliferation in these experimental conditions, confirming that a secreted molecule was causing the death of the phototroph.

### Phototrophs elicit the production of novel cryptic metabolites in *S. tropica*

We analyzed the co-culture supernatants using non-targeted metabolomics to identify the pool of secondary metabolites secreted by *S. tropica* in response to the different phototrophs. The *Synechococcus-S. tropica* co-culture revealed eight molecular ions that were not present in the respective axenic cultures (**Fig. 2, A**). These molecules were further characterized by high-resolution MS/MS analysis, from which we generated empirical chemical formulae, allowing us to assign most of them to two subgroups of related compounds being: (i) ions **1, 2, 5** and **8**; and (ii) ions **4, 6** and **7** (**Fig. 2, A** and **Supplementary Table S1**).

**Figure 2.**
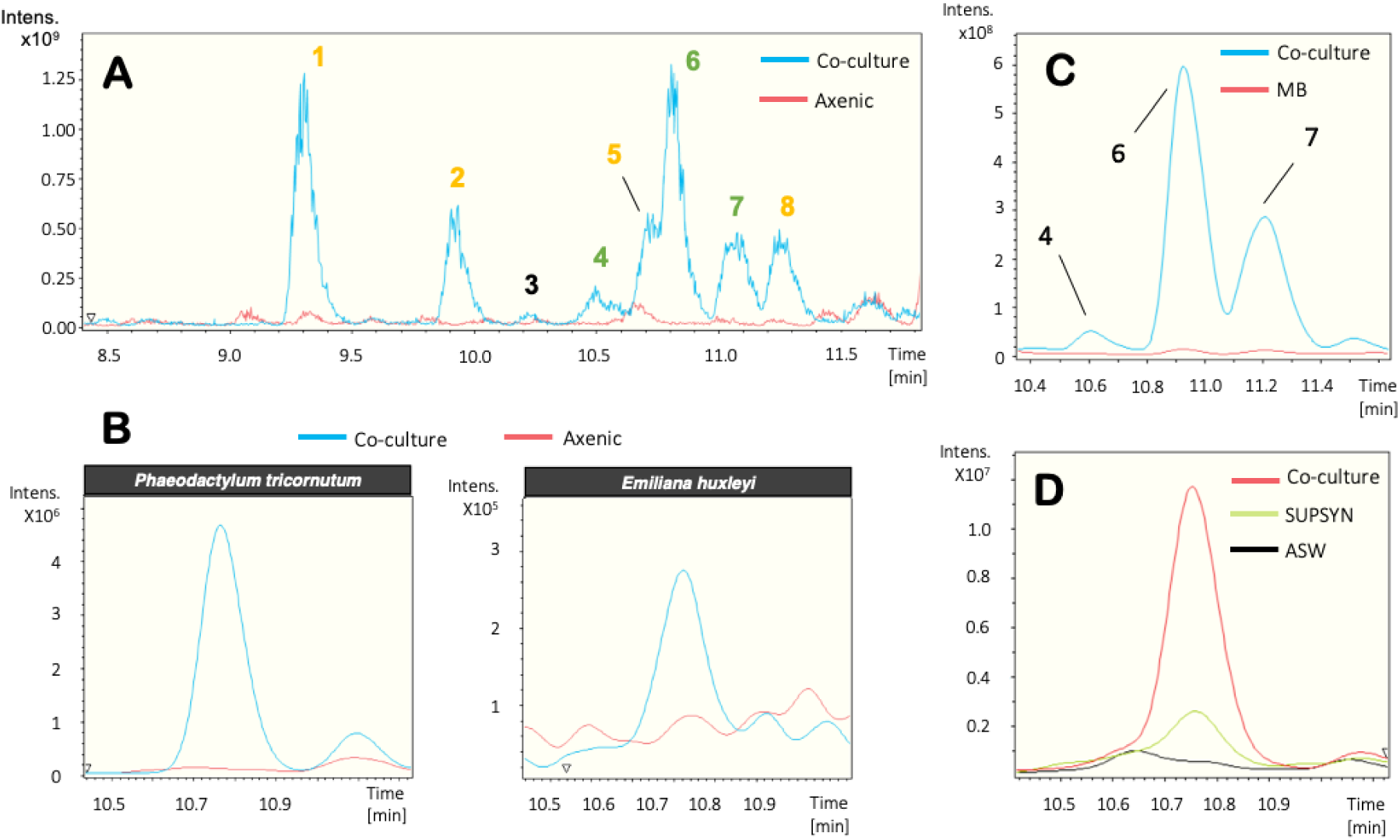
Marine phototrophs trigger the production of cryptic molecules in *S. tropica*. **(A)** *S. tropica* produces detectable small molecules in co-culture with *Synechococcus*. Overlaid base peak chromatograms (BPCs) of *Synechococcus* culture concentrated supernatants, when grown in artificial sea water (ASW) either axenically (red) or in co-culture with *S. tropica* (blue). Peaks characteristic of the co-culture condition are labelled from **1** to **8**. Color of the labels indicate groups of related compounds. **(B)** Other marine phototrophs also trigger the production of metabolite **6** by *S. tropica* as observed in panel A. Figure shows extracted ion chromatograms for the molecule **6** EIC 435.2 ± 0.1 in the supernatants of the phototrophs grown axenically (red) and in co-culture with *S. tropica* (blue). **(C)** The production of the related molecules **4, 6** and **7** is dependent on the presence of photosynthate rather than high-nutrient availability. Graph shows extracted ion chromatograms for all three cryptic molecules EIC (464.2; 435.2; 449.2) ± 0.5 in the concentrated supernatants of *S. tropica* grown axenically in marine broth (MB, red) or in co-culture with *Synechococcus* in ASW (Co-culture, blue). **(D)** Cryptic molecule production is triggered by nutrients released by *Synechococcus* rather than cell-to-cell interactions. Graph shows extracted ion chromatograms for the cryptic molecule **6** (EIC 435.2 ± 0.5) in the supernatant of *S. tropica* grown axenically either in artificial sea water (ASW, black line) or in a conditioned *Synechococcus* supernatant (SUPSYN, green line); and in co-culture with *Synechococcus* (Co-culture, red line).

Ions **1, 2, 5** and **8** were derivatives of salinosporamide; a well-characterized molecule produced by *S. tropica* that presents a unique fused γ-lactam-β-lactone bicyclic ring structure (Feling *et al*., 2003), and that is now being tested as a drug because of its anti-cancer properties. Molecules **5** and **8** are consistent with known degradation products of salinosporamide A and B, respectively (Denora *et al*., 2007; **Supplementary Fig. S2**), while molecules **1** and **2** are proposed to result from the nucleophilic addition of Tris (the buffering agent used in the ASW culture medium) to the lactone ring of salinosporamide A and B, respectively (**Supplementary Fig. S2**). These salinosporamide sub-products were further confirmed by their absence when i) Tris was not added (**Supplementary Fig. S3**), or ii) salinosporamide mutants that no longer produced these metabolites, *i.e.* salA^-^ and salL^-^ (Eustáquio *et al*., 2009), were used (**Supplementary Fig. S4**). In order to test the activity of salinosporamide and its derivatives on the phototrophs, we co-cultured *Synechococcus* with both salinosporamide mutants. Salinosporamide and its derivatives were not responsible for the antimicrobial activity as both deficient mutants were still able to inhibit the phototroph (**Supplementary Fig. S5**).

The second group of ions, *i.e.* peaks **4, 6** and **7**, were also related. Molecule **6** gave a *m/z* value of 435.2609 [M+H]^+^; based on the accuracy of this value and the isotopic pattern the empirical chemical formula C_22_H_35_N_4_O_5_ was predicted by the DataAnalysis software (**Table 2**). The predicted formula for molecule **4** suggests that, with a 28.9900 Da mass difference when compared to **6**, the compound had lost one hydrogen and gained an atom of nitrogen and oxygen. MS/MS analyses confirmed that both molecules **4** and **6** had an identical molecular fragment (*i.e. m/z* 276.1600 ± 0.0001 [M+H]^+^, with the empirical chemical formula C_16_H_22_NO_3_), indicating that the two molecules share a core backbone (**Table 2**). Similarly, molecule **7** had the same chemical formula than **6** but with the addition of a methyl group (14.0155 Da mass difference; **Table 2**). Molecule **3** did not share an obvious link to any other metabolites and, therefore, was considered a new biosynthesized product of *Salinispora* (**Table 2**). Most interestingly, the search for compounds with the same molecular formulae as **3, 4, 6** or **7** in multiple databases (*e.g.* Reaxys, SciFinder, Dictionary of NP) returned no known natural product, suggesting that they are novel compounds. Unfortunately, despite multiple attempts, the isolation of these molecules has so far proven too challenging for their structural elucidation.

**Table 2.**
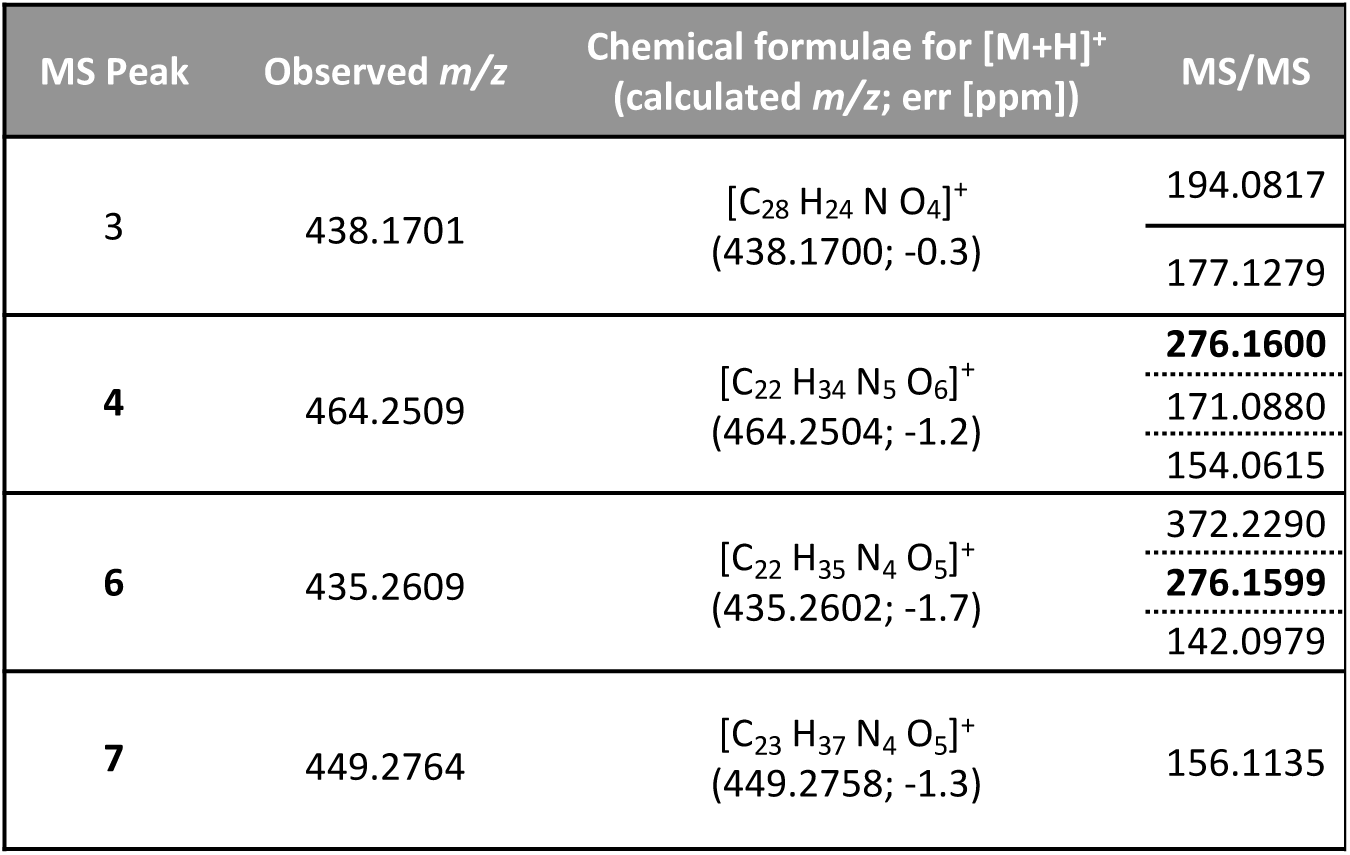
Characteristics of the cryptic molecules. MS Peak numbering is based on HPLC retention time. High-resolution LC- (+)ESI-MS *m/z* values and predicted chemical formulae for [M+H]^+^ are provided.

| MS Peak | Observed *m/z* | Chemical formulae for [M+H]<sup>+</sup><br>(calculated *m/z*; err [ppm]) | MS/MS |
| --- | --- | --- | --- |
| 3 | 438.1701 | [C<sub>28</sub> H<sub>24</sub> N O<sub>4</sub>]<sup>+</sup><br>(438.1700; -0.3) | 194.0817 |
|  |  |  | 177.1279 |
| 4 | 464.2509 | [C<sub>22</sub> H<sub>34</sub> N<sub>5</sub> O<sub>6</sub>]<sup>+</sup><br>(464.2504; -1.2) | **276.1600** |
|  |  |  | 171.0880 |
|  |  |  | 154.0615 |
| 6 | 435.2609 | [C<sub>22</sub> H<sub>35</sub> N<sub>4</sub> O<sub>5</sub>]<sup>+</sup><br>(435.2602; -1.7) | 372.2290 |
|  |  |  | **276.1599** |
|  |  |  | 142.0979 |
| 7 | 449.2764 | [C<sub>23</sub> H<sub>37</sub> N<sub>4</sub> O<sub>5</sub>]<sup>+</sup><br>(449.2758; -1.3) | 156.1135 |

The production of these novel compounds was only triggered by the presence of the phototrophs as they were only detected in the co-cultures of all three phototrophs (**Fig. 2, A and B**), but not when grown in mono culture – as shown by the absence of these metabolites when *S. tropica* was grown alone in mineral ASW or nutrient rich media MB (**Fig. 2, C and D**). Furthermore, we confirmed that the supernatant of a phototroph culture – containing the photosynthate – was enough to induce such metabolite production (**Fig. 2, D**).

### Photosynthate triggers the expression of orphan gene clusters in *S. tropica*

Having detected novel secondary metabolites produced by *S. tropica* in response to phototroph-released photosynthate, we set out to investigate how it affected the induction of its BGCs. To this end, we analyzed and compared the proteome of *S. tropica* when grown in presence of the phytoplankton’s photosynthate – *i.e.* in conditioned *Synechococcus* supernatant – and in nutrient rich broth – *i.e.* marine broth. Surprisingly, we were able to detect proteins encoded by almost all of *S. tropica*’s BGCs, including 10 of its 11 orphans BGCs (**Fig. 3, A**).

**Figure 3.**
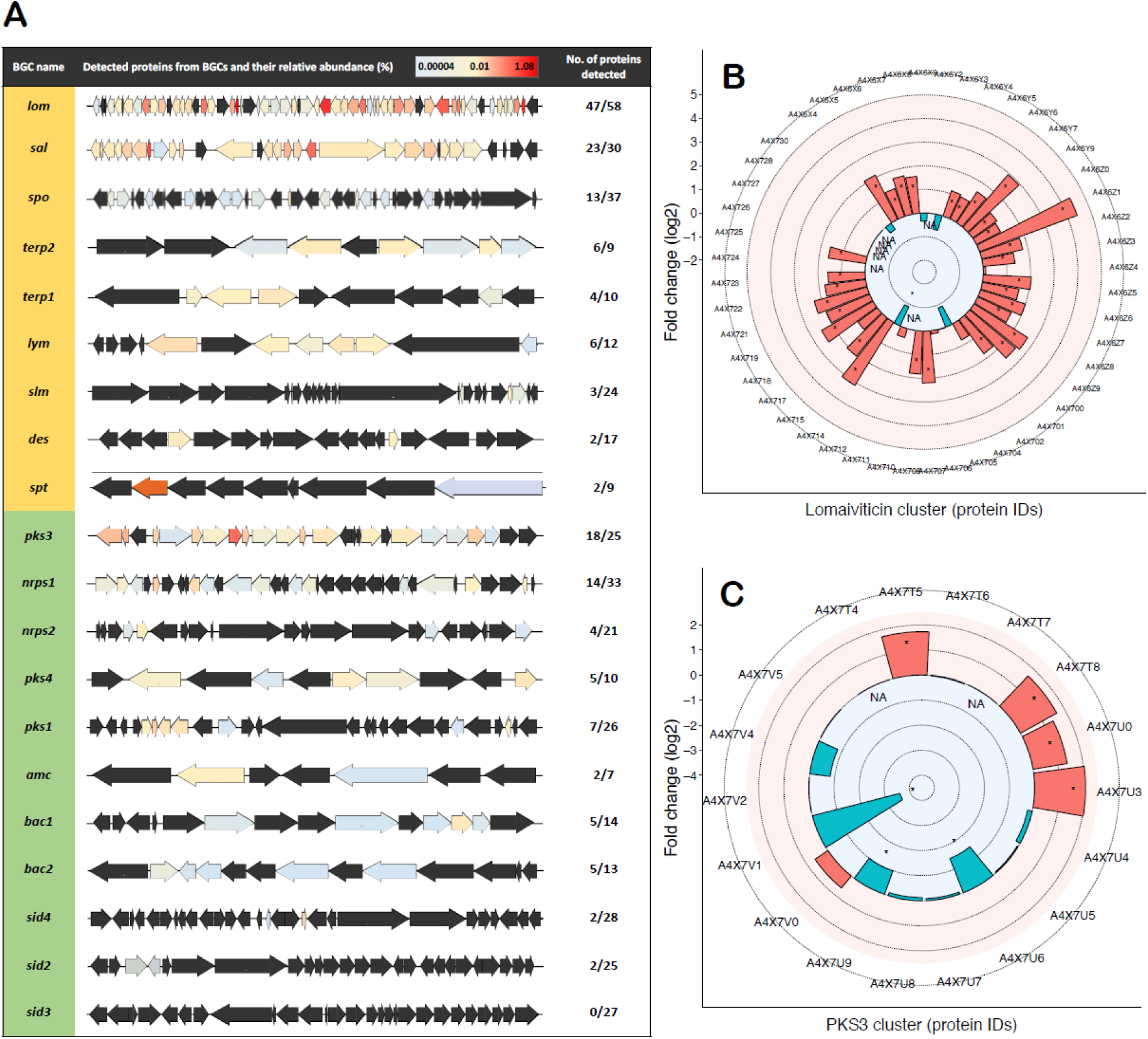
Photosynthate activates orphan biosynthetic gene clusters in *S. tropica*. **(A)** Characterized (orange) and orphan BGCs (green) in *S. tropica* CNB-440 detected by high-throughput proteomics when grown with photosynthate (SUPSYN). Genes are colored according to the relative abundance of their corresponding proteins. Those that were not detected are represented in black. Photosynthate increased the detection of proteins involved in the production of lomaiviticin **(B)** and the orphan PKS3 **(C)** in comparison with cells grown in MB. Up- (red) and down-regulated (blue) proteins in the presence of the photosynthate is shown. Statistically significant fold changes are indicated by an asterisk (T-test, significant at *q*-value < 0.05). *NA* indicate proteins for which the T-test and fold change could not be estimated because of missing values within a set of replicates.

Of particular interest were the orphan BGCs *pks3* and *nrps1*, for which we detected 72% (18/25) and 42% (14/33) of their encoded proteins, respectively (**Fig. 3, A**). Moreover, the *pks3* BGC was noticeably highly detected as eight of its detected proteins showed a relative abundance above 0.1% (**Supplementary Table S2**). While it has been previously suggested that *pks3* may produce a spore pigment polyketide, very little experimental evidence is available in the literature, and the product of *pks3* had not been confirmed (Kersten *et al*., 2013). On the other hand, the non-ribosomal peptide synthetase (NRPS) gene cluster *nrps1* has only been predicted to produce a non-ribosomal dipeptide (Penn *et al*., 2009). Intriguingly, the most abundant proteins detected from this *nrps1* BGC were the non-ribosomal peptide synthetase (A4×2Q0), the condensation domain-containing protein (A4×2R5) and an ATP-dependent Clp protease subunit (A4×2S2), with a relative abundance of 0.004%, 0.001% and 0.121%, respectively (**Table 3**). While the two former are thought to direct the biosynthesis of the non-ribosomal peptide, the later may be involved in conferring resistance to the synthesized antimicrobial compound (Kirstein *et al.*, 2009), as further discussed below.

**Table 3.** Detected proteins from the *nrps1* orphan BGC in *S. tropica* CNB-440 grown with photosynthate. Proteins involved in non-ribosomal peptide biosynthesis and antibiotic self-resistance are highlighted in bold.

| Protein ID | Annotation | Relative abundance<br>(%; n = 3) |
| --- | --- | --- |
| A4X2P5 | RidA family protein | 0.014 |
| A4X2P8 | MFS transporter | 0.011 |
| **A4X2Q0** | **non-ribosomal peptide synthetase** | **0.004** |
| A4X2Q2 | SDR family oxidoreductase | 0.002 |
| A4X2R0 | acyl-CoA dehydrogenase | 0.005 |
| A4X2R1 | acyl-CoA dehydrogenase | 0.001 |
| A4X2R4 | D-alanine--poly(phosphoribitol) ligase | 0.004 |
| **A4X2R5** | **condensation domain-containing protein** | **0.001** |
| A4X2R7 | argininosuccinate synthase | 0.001 |
| A4X2R8 | methionyl-tRNA formyltransferase | 0.087 |
| **A4X2S2** | **ATP-dependent Clp protease proteolytic subunit** | **0.121** |
| A4X2S4 | potassium channel family protein | 0.002 |
| A4X2S5 | 2-oxoacid:ferredoxin oxidoreductase subunit beta | 0.006 |
| A4X2S6 | 2-oxoacid:acceptor oxidoreductase subunit alpha | 0.004 |

The already characterized *lom* and *sal* BGCs were also abundantly detected with 81% (47/58) and 77% (23/30) of their encoded proteins detected, respectively, some representing high relative abundances within the proteome (**Fig. 3, A**). The BGC *lom* is linked to the cytotoxic glycoside lomaiviticin molecule (Kersten *et al*., 2013). However, this metabolite previously showed no antimicrobial activity on co-cultured heterotrophic organisms (Patin *et al.*, 2018) and, hence, it is unlikely to cause the antimicrobial phenotype observed on the phototrophs in this study. The high abundance of the *sal* cluster, producing the salinosporamide compound, is not surprising given the high detection of this metabolite by LC-MS (**Fig. 2, A**).

Interestingly, the comparative proteomic analysis of *S. tropica* grown in photosynthate *versus* marine broth confirmed that the detection of several BGCs rose in response to phototroph-released nutrients, being *lom* and *pks3* the most remarkable ones (**Fig. 3, B-C**). For instance, the *lom* cluster had 77% (36/47) of its detected proteins overexpressed in the presence of the photosynthate (**Fig. 3, B**). The orphan *pks3* BGC was also triggered by the photosynthate, as the pivotal biosynthetic enzymes for polyketide biosynthesis, *i.e.* acyl-CoA ligase (A4×7T8), 3-ketoacyl-ACP synthase (A4×7U0) and long-chain fatty acid-CoA ligase (A4×7U3), were up-regulated (3.1, 2.6 and 4.1-fold change, respectively; **Fig. 3, C** and **Supplementary Table S2**).

## Discussion

We show that *S. tropica* is able to inhibit the growth of both marine cyanobacteria and eukaryotic phototrophs by some, yet, unidentified mechanism (**Fig. 1**). This observation broadens the potential role and impact that the *Salinispora* genus has on marine microbial communities. *Salinispora* is a widely-distributed bacterium found in all tropical and subtropical oceans (Mincer *et al*., 2002; Bauermeister *et al*., 2018). While mostly inhabiting marine sediments, bacteria from this genus were also isolated from marine sponges where it is suggested they influence the sponge microbiota through the production of acyl homoserine lactone molecules and antibiotics (Singh *et al*., 2014; Bose *et al*., 2017). Similarly, different species of *Salinispora* were shown to possess distinct mechanisms to outcompete co-occurring marine heterotrophs in sediments, *i.e.* through the production of siderophores to deplete iron or antimicrobial molecules (Patin *et al*., 2017; Tuttle *et al*., 2019), although no antimicrobial compound has yet been identified for *S. tropica* (Patin *et al.*, 2018). We herein provide the first evidence that *Salinispora* might not only directly influence heterotrophic communities, but also kill both prokaryotic and eukaryotic phytoplankton to which they are exposed, *e.g.* when these sediment out of the water column or phototrophs able to grow in sunlit coastal sediments.

While we were successful in identifying and obtaining the molecular formulae of novel cryptic metabolites produced in response to phytoplanktonic photosynthate (**Fig. 2, Table 2**), we were unable to isolate and identify the compound responsible for the antimicrobial effect on the marine phototrophs by using traditional bioactivity-guided assays with HPLC fractionation of crude extracts (data not shown). This mechanism proved similarly elusive in previous studies, where *S. tropica* showed an antimicrobial activity on marine heterotrophs, but the molecule responsible was not identified (Patin *et al*., 2016; Patin *et al*., 2018). The parallelism between our observations and those described in the literature suggests that the active compound(s) may be the same. We reason that the compound’s instability, and/or synergic effect of several molecules required for activity, could explain the difficulty in identifying the antimicrobial agent. For instance, the large number of structurally-related metabolites resulting from the chemical reaction of salinosporamide with various compounds (*i.e*. water, Tris) may support this hypothesis, as the antimicrobial molecule may be similarly unstable. The diversity of products arising from a single BGC may also be due to the promiscuity of the biosynthetic enzymes utilizing structurally related primary precursors. This results in a range of compounds, each produced at lower titers than a single natural product, and ultimately hamper the isolation of sufficient amounts of the compounds of interest. Whichever the case, we show that *Salinispora* can produce a broad-range antibiotic able to affect both unicellular prokaryotes and eukaryotes alike, such as the marine diatom and coccolithophore tested in our study. Such a broad-range antimicrobial could suggest a mode of action affecting a common target present in both types of cells such as the proteasome, a proteolytic complex present in the three domains of life (Becker and Darwin, 2016).

Exploring the proteome of *S. tropica* exposed to photosynthate, we detected proteins encoded by almost all its BGCs, including most of its orphan BGCs (**Fig. 3**). Notably, the *sal* BGC, producing the salinosporamide compound, was one of the most highly expressed BGC as most of its proteins were detected with high relative abundance. This finding is in agreement with previous studies that have shown by transcriptomics that the BGC *sal* is highly and constitutively expressed when grown in nutrient rich A1 medium (Amos *et al*., 2017). Also, the high expression of this BGC correlated with a noticeable detection of salinosporamide derivatives by LC-MS. The agreement between the metabolomic and proteomic data suggests that it is possible to correlate activated BGCs with the actual biosynthesis of their corresponding natural product. Therefore, the abundant detection of several orphans BGC proteins, including those from *pks3* and *nrps1* BGCs, may be promising candidates responsible for the biosynthesis of the cryptic metabolites detected by LC-MS and, potentially, the antimicrobial activity observed on co-cultured phototrophs.

The proteins detected from the BGC *nrps1* are essential enzymes involved in non-ribosomal peptide biosynthesis, *i.e.* A4×2Q0, a non-ribosomal peptide synthetase (NRPS) made of a C-A-PCP domain, and A4×2RS, a condensation domain-containing protein made of C-PCP-TE domain. The detection of these proteins therefore strongly suggests the actual synthesis of the non-ribosomal peptide and could well be the novel metabolites detected by LC-MS, which include four nitrogen atoms in their predicted molecular formulae (**Table 2**). Interestingly, the substrate specificity of A4×2Q0’s A-domain is alanine and another three A-domains are encoded in the *nrps1* BGC. Further work is required to elucidate the structure of this series of cryptic metabolites. From this same BGC we also detected a highly abundant ATP-dependent Clp protease proteolytic subunit (ClpP, A4×2S2) that may be providing *Salinispora* with self-resistance against the *nrps1* peptides. Virtually all organisms across the tree of life have a system for targeted proteolysis for protein turnover, with most bacteria, mitochondria and chloroplasts relying on a ClpP-type proteasome while eukaryotes, archaea and some actinobacteria typically possess the homologous 20S proteasome structure (Becker and Darwin, 2016; Snoberger *et al*., 2017). The ClpP proteasome is known to be the target for certain antibiotics, including the novel acyldepsipeptide (ADEP) class (Kirstein *et al.*, 2009), and it is common to find alternative ClpP proteasomes encoded nearby the antibiotic-producing BGC to confer resistance to the host cell (Thomy *et al*., 2019). In a similar fashion, salinosporamide A is a 20S proteasome inhibitor, to which *Salinispora* is resistant because of an extra copy of the proteasome beta subunit gene within the salinosporamide-producing cluster (Kale *et al*., 2011). We can thus reasonably infer from the presence of *clpP* in the *nrps1* BGC that it is likely to produce an antibiotic targeting the ClpP proteasome, a class of antimicrobial compounds that has recently gained considerable attention as an attractive option to tackle multidrug resistant pathogens (**Fig. 4**; Momose and Kawada, 2016; Culp and Wright, 2017; Moreno-Cinos *et al.*, 2019). We here provide the first proteomic evidence that *S. tropica*’s *nrps1* is active and may produce a promising antimicrobial compound acting as a ClpP proteasome inhibitor. The synthesis of such antibiotic would explain the antimicrobial effect of *Salinispora* on all marine phototrophs tested in our study as they all rely on the ClpP proteolytic machinery (Andersson *et al.*, 2009; Jones *et al.*, 2013; Zhao *et al.*, 2018). Additional evidence, such as genetic inactivation of the *nrps1* BGC, will confirm this mechanism.

**Figure 4.**
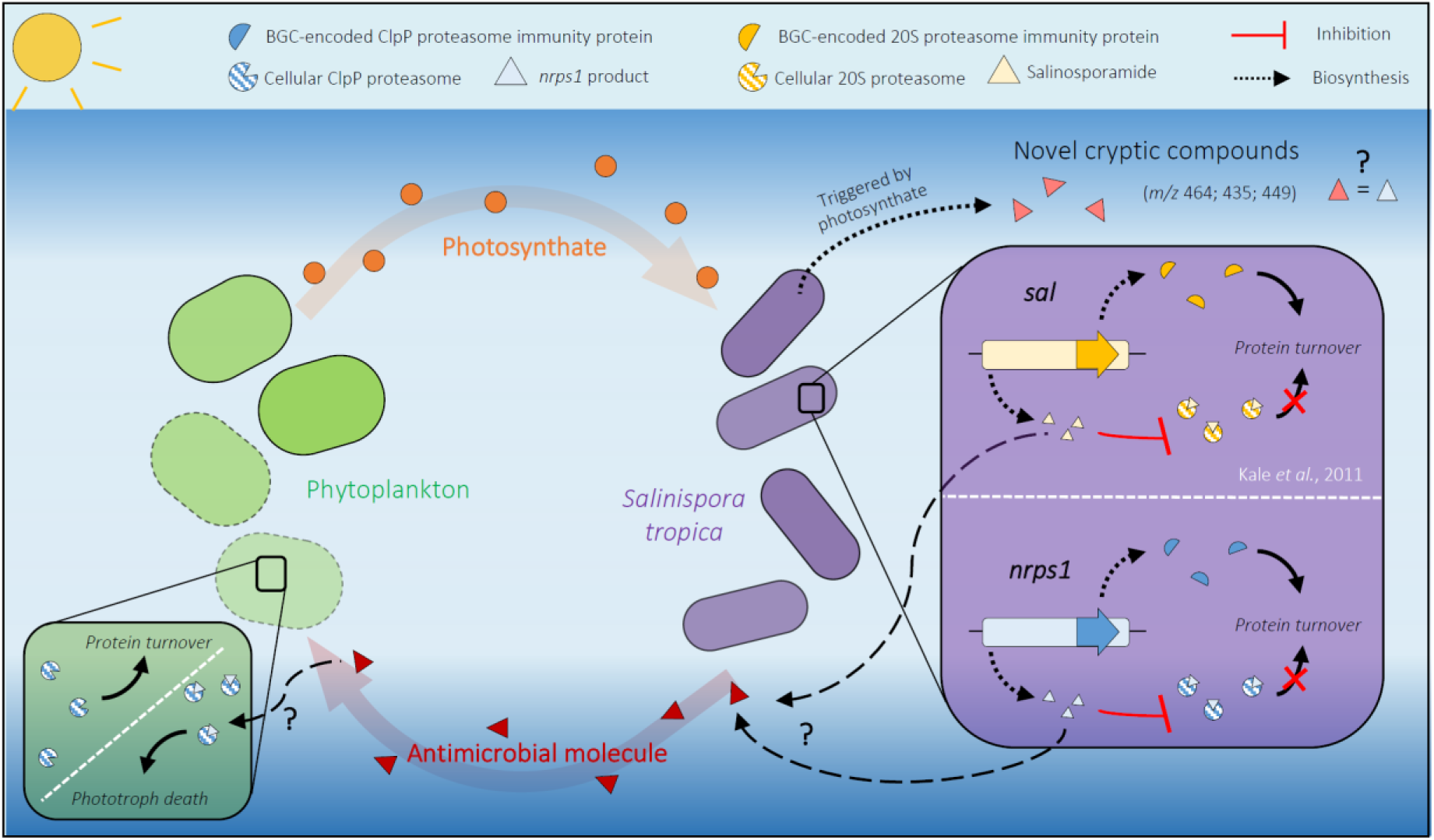
Interaction of *Salinispora tropica* with phytoplankton. Marine phototrophs release photosynthate that triggers the biosynthesis of novel cryptic metabolites in *S. tropica. S. tropica* produces an unknown antimicrobial molecule that kills phytoplankton. The proposed mechanism of the antimicrobial metabolite as well as the activity of the *nrps1* product are depicted (green and purple boxes, respectively). The BGC *nrps1* would produce a ClpP-proteasome inhibitor, to which *S. tropica* would be resistant because of an immunity protein encoded within the BGC, similarly to what is known for *sal*/salinosporamide. The *nrps1*-encoded proteasome inhibitor could kill the phototrophs by preventing protein turnover, leading to cell death.

We show that the photosynthate released by primary producers influences the biosynthetic capacities of *Salinispora*, activating the expression of several orphan BGCs and inducing the production of novel metabolites. Our metabolomics analysis further confirmed the potential of co-culturing for natural product discovery as we identified novel cryptic secondary metabolites, although future work is required to elucidate the structure of the new molecules. Finally, our study extends the pool of known compounds produced by the genus *Salinispora* and pioneers the use of phototrophs as a promising strategy to trigger novel natural products from marine actinobacteria. We also provide a valuable insight into the biosynthetic potential of *S. tropica* with our proteomic dataset, which reveals the *nrps1* BGC as a promising candidate for antibiotic production.

## Supporting information

Supplemental File 1 (proteomic dataset)

## Acknowledgments

We thank Vinko Zadjelovic, Linda Westermann and Fabrizio Alberti for helpful discussions throughout the project. We also acknowledge technical support from Cleidiane Zampronio of the WPH Proteomic Facility at the University of Warwick. In addition, we thank the BBSRC/EPSRC Synthetic Biology Research Centre WISB BB/M017982/1 for access to the flow cytometer and Yin Chen for access to the LC-MS. A.C. was supported by an MIBTP PhD scholarship (BB/M01116X/1) and D.S. by a NERC CENTA DTP studentship (NE/L002493/1). J.A.C.-O was funded by a NERC Independent Research Fellowship NE/K009044/1 and Ramón y Cajal contract RYC-2017-22452 (funded by the Ministry of Science, Innovation and Universities, the National Agency of Research, and the European Social Fund). C.C. thanks BBSRC (grant BB/M022765/1) and European Union’s Horizon 2020 research No. 765147 for support. L.S. would like to acknowledge BBSRC (BB/M017982/1 and BB/R000689/1) and EPSRC (EP/P0305721/1) for financial support.

