## Supplemental Figures S1-S5 and Tables S1-S2 for "Phytoplankton trigger the production of cryptic metabolites in the marine actinobacteria *Salinispora tropica*"

---

### Supplementary Figures

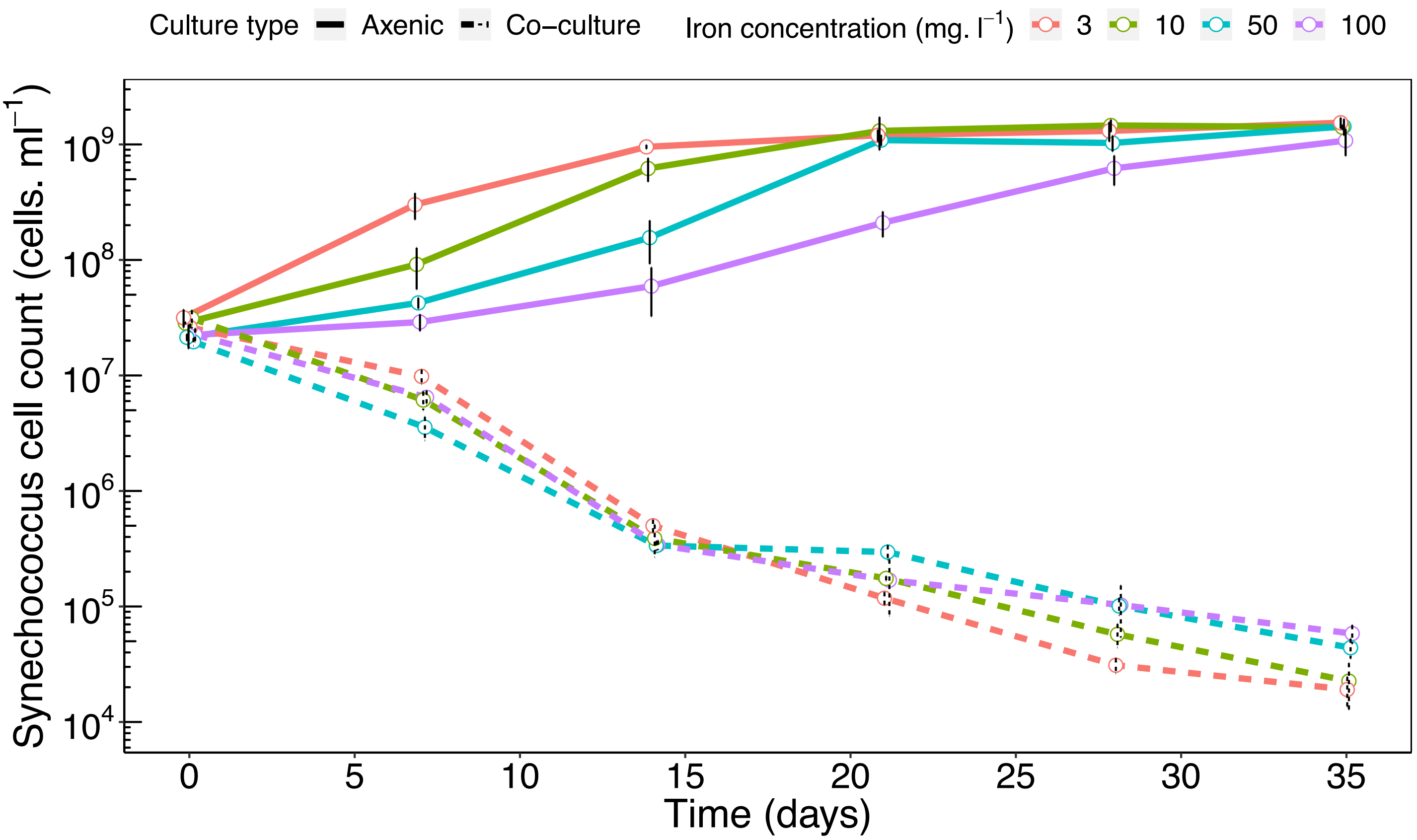


**Supplementary Figure S1 | *Synechococcus* inhibition by *S. tropica* is not mediated by iron depletion**. Monitoring of *Synechococcus* population grown axenically or in co-culture with *S. tropica*, in media supplemented with 3, 10, 50 or 100 mg $l^{-1}$ Fe(III). Graph shows mean ± standard deviation of three biological replicates.

**Supplementary Table S1 | Molecular ions detected by LC-MS in *S. tropica-Synechococcus* co-culture only.** Table shows molecular ions detected by high-resolution LC/(+)ESI-MS. Peak numbering is based on HPLC retention time and colors indicate groups of related compounds. Observed *m/z* values and predicted chemical formulae for $[M+H]^+$ are provided. Observed mass of main ions obtained after MS2 fragmentation are given.

| MS Peak | Observed *m/z* | Chemical formulae for $[M+H]^+$ (calculated *m/z*; err [ppm]) | MS/MS |
|---|---|---|---|
| 1 | 399.2135 | $[C_{19} H_{31} N_2 O_7]^+$ (399.2126; -2.4) | **296.1498**<br>271.1295<br>186.0765<br>168.0659 |
| 2 | 401.2294 | $[C_{19} H_{33} N_2 O_7]^+$ (401.2292; -2.8) | **298.1657**<br>273.1453<br>255.1348<br>152.0712 |
| 3 | 438.1701 | $[C_{28} H_{24} N O_4]^+$ (438.1700; -0.3) | 194.0817<br>177.1279 |
| 4 | 464.2509 | $[C_{22} H_{34} N_5 O_6]^+$ (464.2504; -1.2) | **276.16**<br>171.088<br>154.0615 |
| 5 | 296.1499 | $[C_{15} H_{22} N O_5]^+$ (296.1492; -2.1) | 296.1499<br>318.132<br>168.066 |
| 6 | 435.2609 | $[C_{22} H_{35} N_4 O_5]^+$ (435.2602; -1.7) | 372.229<br>**276.1599**<br>142.0979 |
| 7 | 449.2764 | $[C_{23} H_{37} N_4 O_5]^+$ (449.2758; -1.3) | 156.1135 |
| 8 | 298.1653 | $[C_{15} H_{24} N O_5]^+$ (298.1649; -1.5) | 320.1447<br>298.1654<br>170.0815 |

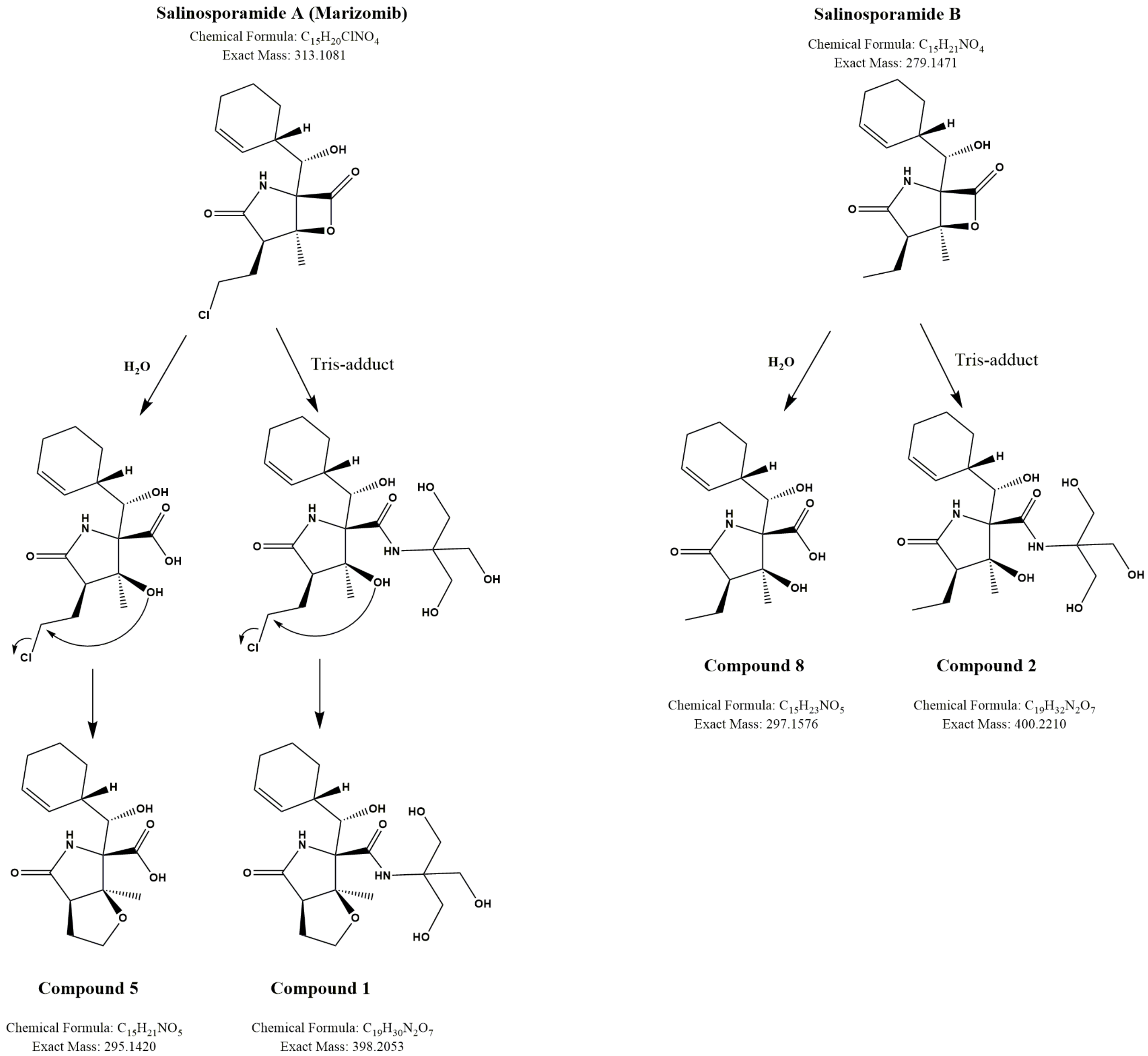


**Supplementary Figure S2 | Chemical structure of salinosporamide A and B, with their respective degradation products.** Salinosporamide A ($C_{15}H_{20}{}^{35}ClNO_4$; mass 313.11) hydrolyzes to form the molecule NPI-0065 (**5**) ($C_{15}H_{21}NO_5$; mass 295.14) or reacts with Tris to form the hypothetical molecule (**1**) ($C_{19}H_{30}N_2O_7$; mass 398.21). Salinosporamide B ($C_{15}H_{21}NO_4$; mass 279.15) hydrolyzes to form the molecule (**8**) ($C_{15}H_{23}NO_5$; mass 297.16) or reacts with Tris to form the hypothetical molecule (**2**) ($C_{19}H_{32}N_2O_7$; mass 400.22).

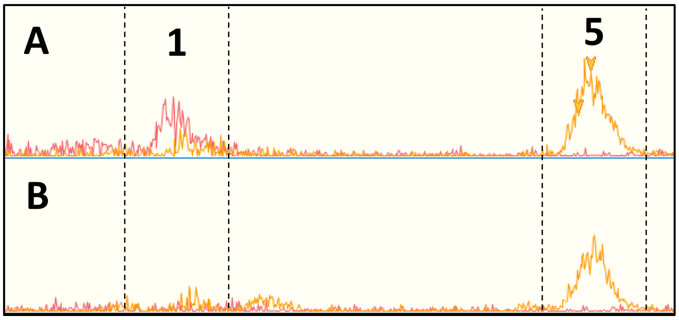


**Supplementary Figure S3 | Extracted Ion chromatograms of molecules 1 and 5 in the supernatant of *S. tropica* cultures in marine broth. A.** Culture supernatant of *S. tropica* grown in marine broth supplemented with trizma base. **B.** Culture supernatant of *S. tropica* grown in marine broth. Graphs show molecules detected with a retention time between 8.8 and 11.1 minutes. In red is shown the extracted ion chromatogram for *m/z* 399 (± 0.5). In orange is shown the extracted ion chromatogram for *m/z* 296 (± 0.5).

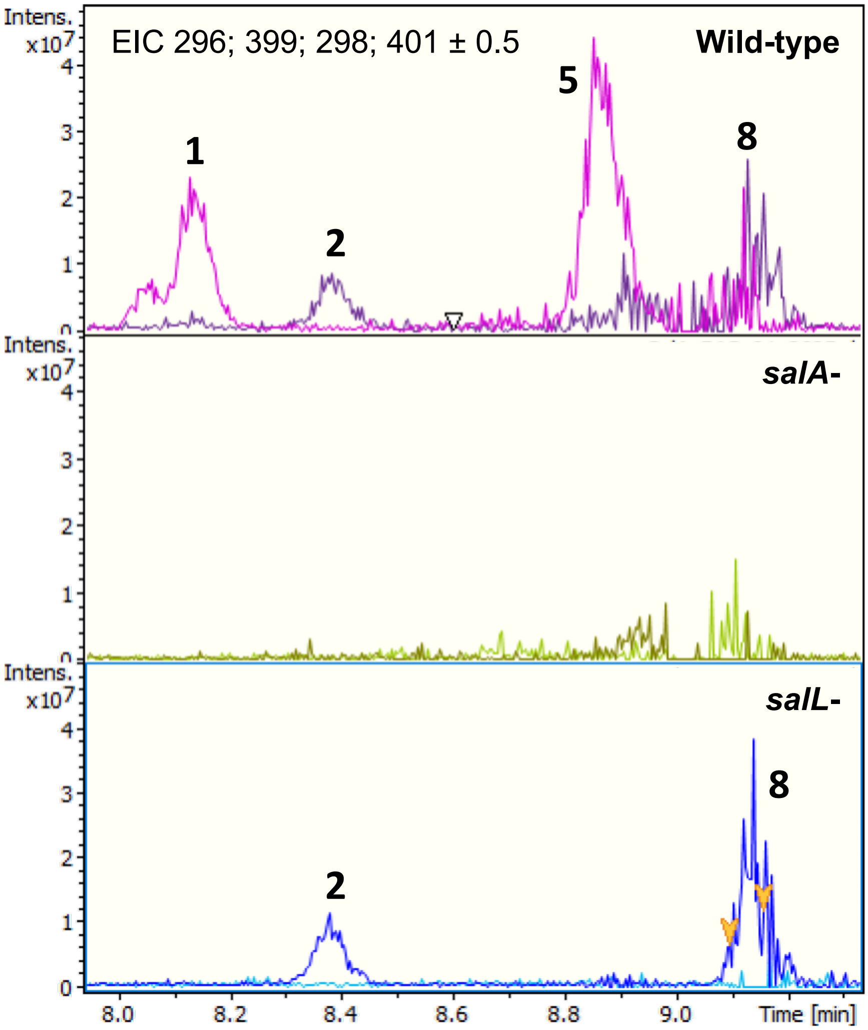


**Supplementary Figure S4 | Extracted ion chromatograms of molecules 1, 2, 5 and 8 in the culture supernatant of *S. tropica* wild-type (top panel), and the salinosporamide mutants *salA*- (middle panel) and *salL*- (bottom panel).** The *salA*- strain does not produce salinosporamide A or any derivatives, while the *salL*- strain still produces salinosporamide B. Graphs show molecules detected with a retention time between 8.0 and 9.4 minutes.

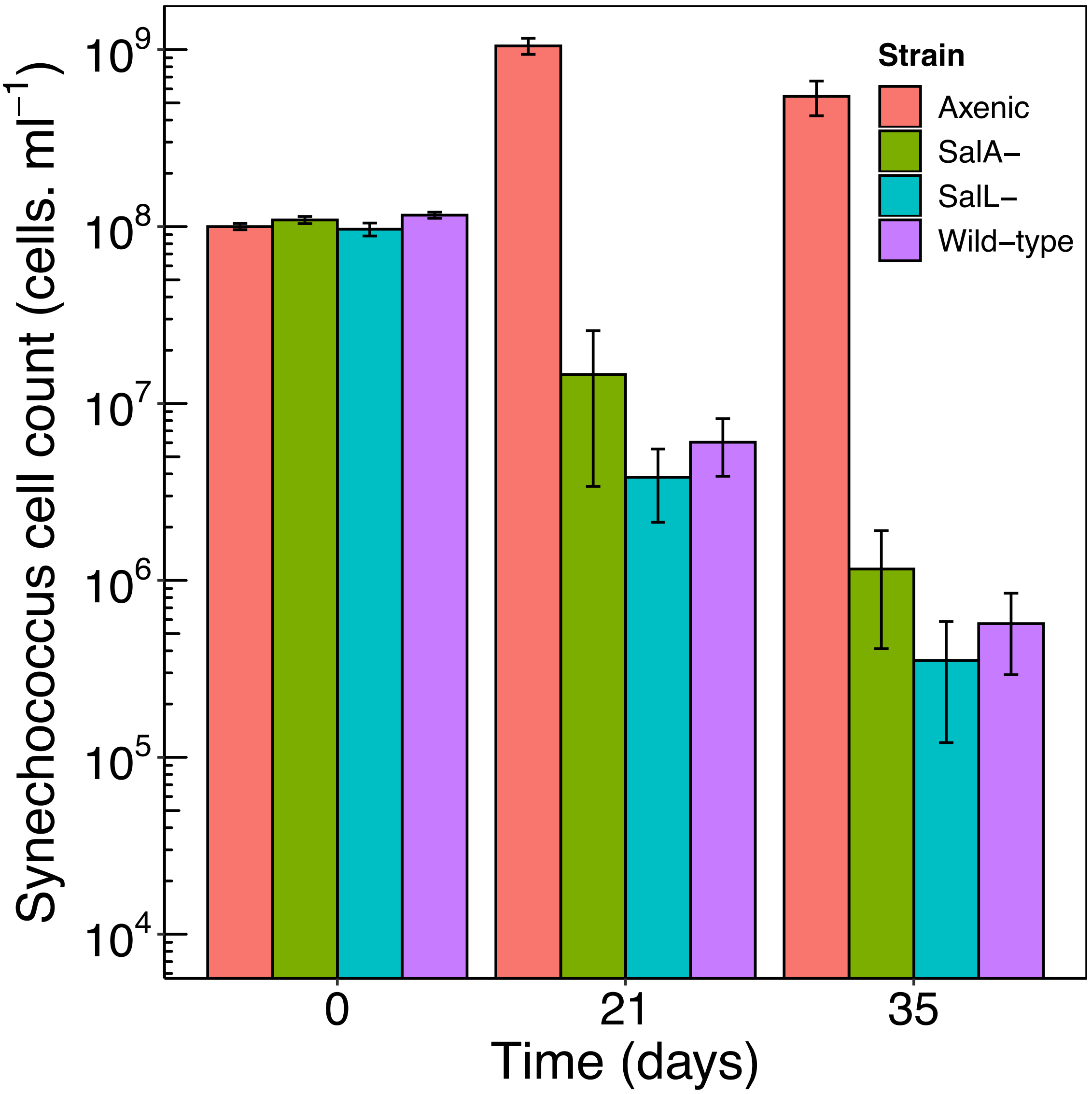


**Supplementary Figure S5 | Monitoring of *Synechococcus* grown in axenic culture and in co-culture with the wild-type, *salA*- or *salL*- *S. tropica* strains.** Graph shows mean of triplicates ± standard deviation.

**Supplementary Table S2 | Detected proteins from the *pks3* orphan BGC in *S. tropica* CNB-440.** Table shows protein identifiers, annotation and relative abundance (expressed as the abundance of the protein over the abundance of the total proteome normalized to 1).

| Protein ID | Gene name | Locus tag | Annotation | Relative abundance |
|---|---|---|---|---|
| A4X7T4 | Strop_2488 | STROP_RS12520 | DUF3050 domain-containing protein | 0.000750547 |
| A4X7T5 | Strop_2489 | STROP_RS12525 | phytanoyl-CoA dioxygenase | 0.130375917 |
| A4X7T6 | Strop_2490 | STROP_RS12530 | 4-hydroxyphenylpyruvate dioxygenase | 0.002402502 |
| A4X7T7 | Strop_2491 | STROP_RS12535 | beta-ketoacyl | 0.001818263 |
| A4X7T8 | Strop_2492 | STROP_RS12540 | acyl--CoA ligase | 0.031677582 |
| A4X7U0 | Strop_2494 | STROP_RS12550 | 3-ketoacyl-ACP synthase | 0.014597734 |
| A4X7U3 | Strop_2497 | STROP_RS12570 | long-chain fatty acid--CoA ligase | 0.034956859 |
| A4X7U4 | Strop_2498 | STROP_RS12575 | antibiotic biosynthesis monooxygenase | 0.129801561 |
| A4X7U5 | Strop_2499 | STROP_RS12580 | actinorhodin polyketide beta-ketoacyl synthase | 0.006074535 |
| A4X7U6 | Strop_2500 | STROP_RS12585 | beta-ketoacyl | 0.00502919 |
| A4X7U7 | Strop_2501 | STROP_RS12590 | cupin domain-containing protein | 0.146326909 |
| A4X7U8 | Strop_2502 | STROP_RS12595 | cyclase | 0.495965212 |
| A4X7U9 | Strop_2503 | STROP_RS12600 | acetyl-CoA carboxylase biotin carboxylase subunit | 0.006262552 |
| A4X7V0 | Strop_2504 | STROP_RS12605 | acetyl-CoA carboxylase biotin carboxyl carrier protein | 0.10444248 |
| A4X7V1 | Strop_2505 | STROP_RS12610 | acetyl-CoA carboxylase carboxyltransferase subunit alpha | 0.000773389 |
| A4X7V2 | Strop_2506 | STROP_RS12615 | TcmI family type II polyketide cyclase | 0.117717019 |
| A4X7V4 | Strop_2508 | STROP_RS12625 | TcmI family type II polyketide cyclase | 0.229200995 |
